# Inflammation-driven reprogramming of goblet cells underlies the onset of serrated adenomas in colon cancer

**DOI:** 10.64898/2026.06.22.733748

**Authors:** Mathijs P. Verhagen, Emily Middendorp Guerra, Rosalie Joosten, Andrea Sacchetti, Wenjie Sun, Karishma Lila, Alex Nigg, Eiman I. Ahmed, Heba Saadeh, Thierry P.P. van den Bosch, Wouter Hendrickx, Jessica Roelands, Annemarie C. de Vries, Michail Doukas, Riccardo Fodde

**Affiliations:** Department of Pathology, Erasmus University Medical Center, Rotterdam, The Netherlands; Institut Curie, Laboratory of Genetics and Developmental Biology, Paris, France; Translational Medicine Division, Research Branch, Sidra Medicine, Doha, Qatar; Department of Computer Science, KASIT, The University of Jordan, Amman, Jordan; Department of Pathology, Leiden University Medical Center, Leiden, The Netherlands; Department of Gastroenterology and Hepatology, Erasmus University Medical Center, Rotterdam, The Netherlands

**Author notes:** Department of Discovery Oncology, GenenTech, South San Francisco, CA, U.S.A. Institute for Research in Biomedicine (IRB Barcelona), The Barcelona Institute of Science and Technology (BIST), Barcelona, Spain. van Andel Institute, Grand Rapids, MI, U.S.A.

## Abstract

In the context of inflammation, fully committed and post-mitotic cell lineages can initiate intestinal tumorigenesis in the mouse through dedifferentiation and acquisition of revival stem cell (RSC) features^1,2^. Likewise, by means of machine-learning analysis of whole-genome mutation spectra, the secretory goblet cell was predicted as the most frequent cell-of-origin (COO) of colon cancer in inflammatory bowel disease (IBD) patients^2^. Of note, even among sporadic (non-IBD) colon cancer patients, goblet cells were predicted as the most common differentiated COO of cancer in the colonic epithelium^2^, suggestive of the main role played by diet-induced inflammation in the onset of a large proportion of malignancies of the large bowel. However, how goblet cells respond to inflammation and reprogram their identity to become potential tumor-initiating cells remains unclear.

Here, by taking advantage of publicly available single-cell RNAseq data from colonic tissues of ulcerative colitis (UC) patients^3^, we have characterized the inflammation-driven reprogramming of goblet cells. By means of an RNA velocity-based computational approach, we show that the mucus-producing goblet cells acquire an aberrant proliferative state earmarked by *MUC5AC*^+^ expression. Trajectory analysis of serrated adenoma cells^4^ reveal aberrant goblet cells as an intermediate state in the transition to revival stem cells, notably more common in *BRAF*-mutant cases. In support of these findings, COO predictions using whole-genome mutation spectra from a cohort of sporadic colon cancers^5^ connect tumors with a predicted goblet origin to *BRAF* mutations and reveal transcriptional remnants of the aberrant goblet cell state.

## Introduction

*Lgr5*^+^ intestinal stem cells (ISCs), also known as crypt-base columnar (CBC) cells, have been shown to represent the obligate origin of intestinal adenomas in the mouse^6^, in support of the current view according to which most tumors arise from stem or progenitor cells^7,8^. However, in the context of inflammation, i.e. one of the main colon cancer risk factors^9^, the lineage tracing capacity of *Lgr5*^+^ ISCs is largely compromised, while more committed lineages, including secretory precursors^10^, tuft^11^, and Paneth cells^2^, are reprogrammed to acquire stem-like features, thus unlocking their tumor-initiating capacity^1^. The first suggestion of this alternative ‘top-down’ route, where differentiated cells represent the origin from which colonic adenomas arise, was provided by Shih et al.^12^ who identified *APC* genetic alterations in dysplastic cells located at the luminal surface of the crypts.

More recently, single-cell RNA sequencing (scRNAseq) analysis of precursor benign lesions also suggested distinct routes through which adenomas may arise^4^. While conventional adenomas appear to result from the expansion of the ISC pool, serrated polyps, earmarked by their saw-tooth histological appearance and strongly associated with increased colorectal cancer (CRC) risk^13^, were suggested to originate from non-stem lineages, possibly as the result of metaplasia-associated reprogramming^4^.

Among the differentiated lineages characteristic of the human colonic epithelium, enterocytes and secretory goblet cells are most abundant. Of note, by means of machine-learning analysis of whole-genome mutation spectra, goblet cells were predicted as the most common non-stem lineage of origin both in IBD-associated and in microsatellite-stable (MSS) sporadic colon cancer^2^.

Evidence from mouse models has shown that, upon irradiation, intestinal tissue regeneration is earmarked by the acquisition of the revival/regenerative stem cell (RSC) state^14^. The same RSC state was observed in dedifferentiating and tumor-initiating secretory cells in the context of inflammation^2^, and has been associated with a distinct subtype of human colon cancers characterized by fetal transcriptional profiles^15^, and with those with a predicted goblet cell origin^2^. However, the mechanisms underlying the inflammation-driven dedifferentiation and reprogramming of fully committed lineages and the (epi)genetic perturbations that predispose these cells to malignant transformation, remain to date unresolved.

Here, we utilize computational methodologies to investigate inflammation-induced reprogramming in the human colon. To this end, we employed scRNAseq data from colonic tissues of healthy donors and ulcerative colitis (UC) patients^3^ and analyzed them by RNA velocity, a method that uses the ratio between spliced and un-spliced mRNA to predict individual cell fate decisions^16,17^. By comparative analysis of the differentiation process in healthy and colitis subjects, we identify a subpopulation of cells participating in deviant differentiation processes. These colitis-induced “aberrant” goblet cells are earmarked by the expression of the otherwise gastric-specific mucin 5AC (*MUC5AC*) gene. Using trajectory analysis of data from pre-malignant polyps^4^, we show that aberrant goblet cells are also present in serrated adenomas, where they represent an intermediate state in the conversion from goblet cells to RSCs. We propose that colitis induces dedifferentiation and aberrant goblet cell formation and thereby enlarges the pool of cell targets for serrated adenoma initiation and progression towards malignancy.

## Results

### RNA velocity analysis identifies aberrant goblet cells in ulcerative colitis

To investigate colitis-induced transcriptional reprogramming, we took advantage of the single-cell atlas previously generated by Smillie et al.^3^ encompassing 283,015 cells derived from the colonic mucosa of 17 UC patients and 12 healthy individuals (Figure 1A, Suppl. Figure 1A). Most UC patients had matched samples from both non-inflamed and inflamed areas, as determined by pathological assessment. Due to the high transcriptional similarity between epithelial cells from matched non-inflamed and inflamed samples, as also reported in the original study, we combined these samples and referred to them collectively as UC.

**Figure 1.**
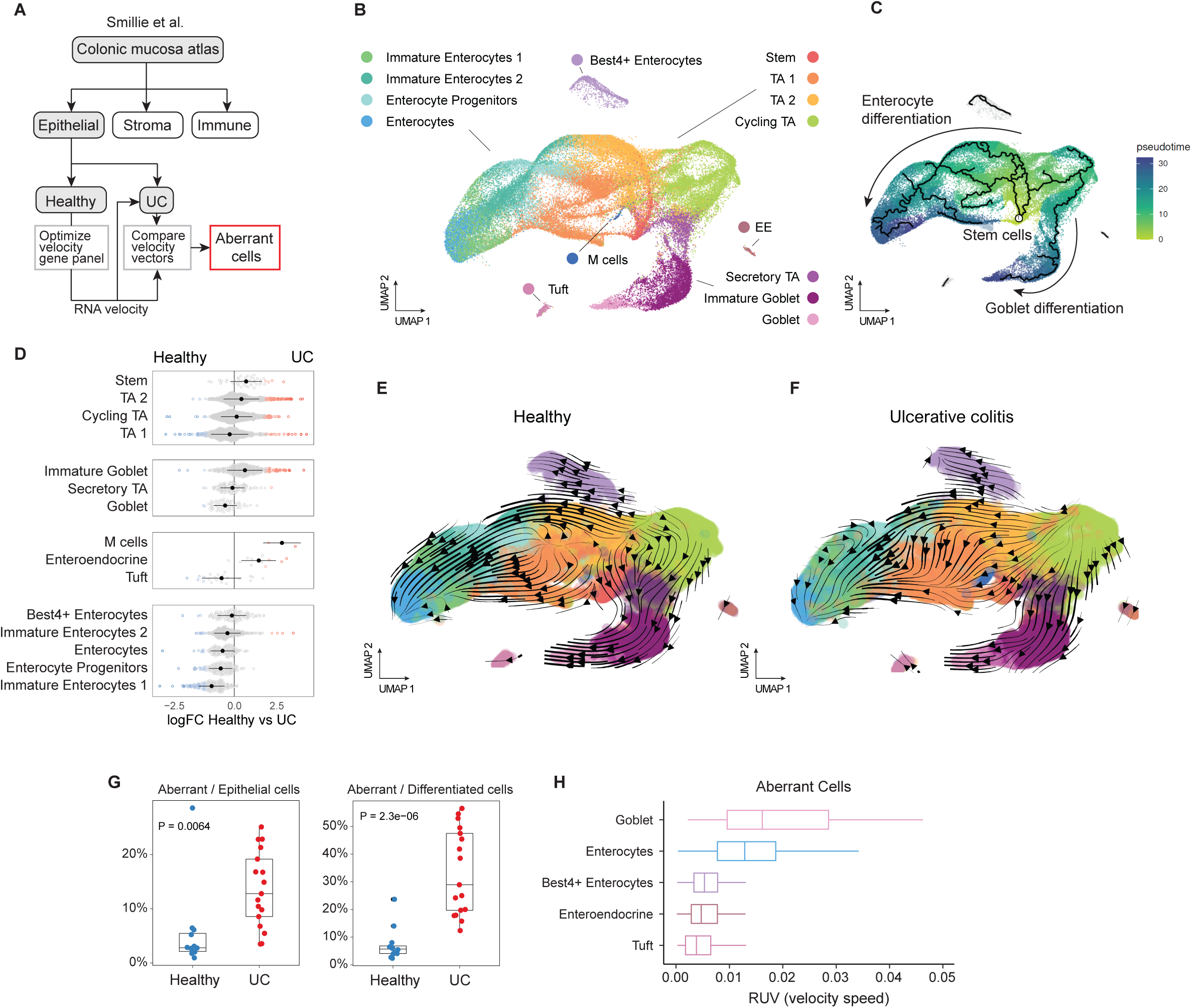
RNA velocity based identification of aberrant cells in ulcerative colitis. **A.** Schematic flowchart describing the pipeline employed for the identification of aberrant cells. **B**. Uniform Manifold Approximation and Projection for Dimension Reduction (UMAP) embedding showing the epithelial cells from the Smillie et al.^3^ data set. TA: transient-amplifying cells. **C**. Feature plot highlighting differentiation trajectories based on pseudotime ordering with Monocle3^48^. **D**. Bee swarm plot showing the results of the differential abundance analysis using Milo^19^. Dots denote neighborhoods that are associated with ulcerative colitis (red) or control samples (blue). Neighborhoods with an FDR > 0.1 are depicted in grey. **E-F**. RNA velocity stream plots produced by scVelo^18^ on the data sets from healthy (**E**), and UC (**F**) patients. **G**. Boxplots showing the percentage of aberrant cells per patient among all epithelial cells (left) and among the differentiated cells (right). Differentiated cells included the annotations of enterocytes, goblet cells, enteroendocrine cells, best4+ enterocytes, and Tuft cells. **H**. Boxplots denoting the distribution of velocity speeds in aberrant cells, separated by annotation.

We employed the RNA velocity tool^17,18^ to identify committed cells predicted to deviate from their normal differentiation trajectories, hereafter referred to as “aberrant” cells (Figure 1A).

Focusing on epithelial cells, we filtered high-quality cells and sampling (see Methods), which resulted in a final dataset of 56,363 epithelial cells (n=28,498 from healthy controls and n=27,865 from UC patients). To visualize the distinct cell lineages throughout the data set, we performed dimension reduction by Uniform Manifold Approximation and Projection (UMAP). After batch correction and the integration of cells from different individuals, the UMAP embedding separated the distinct cell lineages of the colonic epithelium (Figure 1B). Trajectory inference analysis revealed branching of intestinal stem cells into the major absorptive and secretory lineages, with pseudo-time ordering supporting enterocytes and goblet cells as the endpoints of these differentiation trajectories, respectively (Figure 1C). Notably, Best4+ enterocytes, tuft, and enteroendocrine cells formed compact, distinct clusters, separated from the bulk of the other epithelial lineages.

To determine whether UC alters the abundance of certain cell types, we first conducted a differential abundance test by comparing overlapping cellular neighborhoods using Milo (see Methods)^19^. This approach revealed an overall decrease of differentiated neighborhoods in UC, especially notable in enterocytes, goblet, and tuft cells (Figure 1D). Conversely, stem and progenitor cells showed increased abundance in UC, as well as immature goblet, enteroendocrine, and M cells. Of note, the inflammation-driven induction of colonic M and enteroendocrine cells has been previously reported both in IBD patients and in animal models of colitis^20,21^.

Next, to explore if reprogramming of differentiated cells could contribute to this compositional imbalance, we investigated cellular plasticity by RNA velocity modeling. RNA velocity infers directionality of differentiation based on the ratio of spliced to unspliced mRNA molecules^17,18^. We first optimized the use of RNA velocity for this data set by refining the panel of genes that serve as input to the model (see Methods and Suppl. Figure 1B-D). To this end, we selected the velocity genes detected by scVelo and grouped them in three clusters based on their expression patterns in the colonic epithelium (Suppl. Figure 1C). We then manually assessed scatter plots illustrating the relative fraction of spliced vs. un-spliced reads, also referred to as *phase portraits*, for each velocity gene group. During this inspection, we identified one cluster with noisy, cloud-like phase portraits, leading us to exclude it from the set of velocity genes employed in the model (Cluster 2, Suppl. Figure 1D). Of note, Cluster 2 also exhibited lower overall gene counts, suggesting sparse detection of these genes in our dataset.

Using the filtered set of velocity genes (N = 834), we modeled differentiation with a dynamical RNA velocity model using the epithelial cells from healthy individuals. The resulting velocity projection was in agreement with the pseudotime analysis (Figure 1C) and with established biological knowledge, showing the progression from stem cells and transit-amplifying cells to differentiated lineages (Figure 1E). Importantly, applying the same velocity gene panel to an independent dataset^22^ encompassing cells from the small intestine of Crohn’s disease patients produced comparable results, generalizing the applicability of this gene panel for modeling intestinal differentiation (Suppl. Figure 2).

We then employed the same gene panel to run RNA velocity on cells from UC patients (Figure 1F). Projection of the RNA velocity flow on the UMAP embedding revealed comparable results to those obtained from normal samples, suggesting that the gene panel robustly captures overall differentiation dynamics. To evaluate whether specific cell types escape the normal flow of differentiation, we compared the velocity vectors of each cell. To this end, we used a two-step approach. First, we calculated the average angle and standard deviation of differentiation for each cell lineage within the set of cells from healthy individuals. Then, cells were identified as ‘aberrant’ if their velocity angle differed by more than two standard deviations from the average angle. Following this approach, 10.6% of epithelial cells were labeled as aberrant and were significantly more abundant in tissues from UC patients compared to those from healthy subjects, especially within the pool of differentiated cells (Figure 1G).

It should be acknowledged that the utility of this approach for the identification of aberrant cells is facilitated when dealing with differentiated lineages, as they represent the endpoint of a differentiation trajectory. In contrast, stem or progenitor cells can branch off into multiple lineages, which complicates the definition and comparison with the consensus angle. Hence, for the remaining analysis, we focused on aberrant cells from differentiated lineages. Apart from the angle, the RNA velocity model also provides each cell with a velocity length, as a proxy of the predicted speed at which transitions occur. Aberrant cells from differentiated lineages typically displayed low velocity, consistent with their position as the terminal state in the differentiation process. When comparing the length of velocity vectors across aberrant cells from different lineages, aberrant goblet cells displayed the largest vectors, followed by aberrant enterocytes, suggesting that these lineages undergo the most reprogramming in response to inflammation (Figure 1H).

We computed single-cell entropy (SCENT)^23^ to assign to each cell a score of its differentiation potential. As expected, stem and progenitor cells exhibited higher entropy scores compared to differentiated cell lineages (Suppl. Figure 1E). Interestingly, cells from UC patients showed higher entropy values than their healthy counterparts, independent of differentiation status, presumably indicative of their elevated plastic potential (Suppl. Figure 1F).

Taken together, these analyses suggest that inflammation interferes with epithelial differentiation in the colon by triggering aberrant cell-fate trajectories, particularly notable in differentiated cells of the goblet and enterocyte cell lineages.

### Characterization of the aberrant goblet cell state

To explore the inflammation-driven transcriptional alterations associated with aberrant differentiation and reprogramming of the secretory goblet cells, we first visualized their individual velocity vectors in a compass-like plot (Figure 2A). In UC, the fraction of goblet cells with aberrant increased from 1.7 ± 4.2% among healthy subjects to 27.3 ± 32.7% in UC (Figure 2A). To characterize differential gene expression among aberrant cells as defined by RNA velocity analysis, we explored their individual scRNAseq profiles, independent of splicing, by utilizing gene signatures specific for mature and immature goblet cells, as identified in the original Smillie et al. study^3^. While the evaluation of expression signatures of mature and immature goblet cells in healthy colonic tissues revealed two clearly distinguishable states (Figure 2B, left panel), aberrant goblet cells from UC-derived samples exhibited intermediate states between the fully differentiated and more immature cell identities (Figure 2B, right panel) suggestive of a hybrid profile, as also supported by comparative analysis of goblet-specific gene markers: aberrant cells co-expressed markers of mature (*MUC2*, *ZG16*) and immature (*SPINK4*, *SPINK1*) goblet cells (Suppl. Figure 3A).

**Figure 2.**
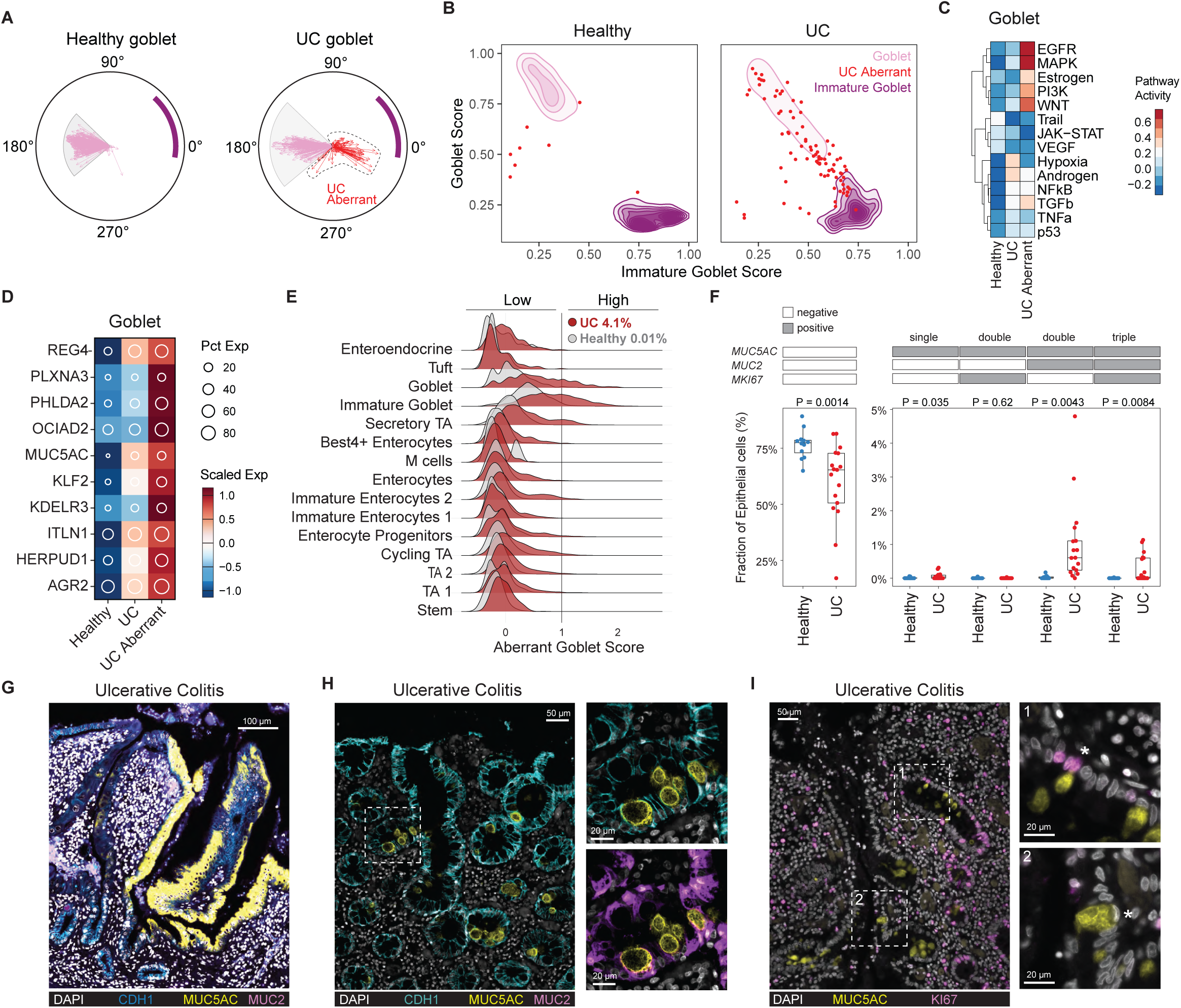
Characterization of colitis-induced aberrant goblet cells. **A.** Compass plot showing the angle and length of the single cell velocity vectors from the goblet cell lineage. The purple bar denotes the position of the immature goblet cells for reference. Separate plots were produced for the healthy (left) and ulcerative colitis (right) data sets. Red arrows indicate aberrant goblet cells. **B**. Density plot showing the distribution of the goblet and immature goblet signatures across the goblet (pink) and immature goblet cells (purple). Aberrant goblet cells (red) were plotted on top as single dots (cells) to highlight their position. **C**. Heat map showing pathway activities across the distinct goblet cell groups. Pathway scores were computed with decoupleR^24^ **D**. Heat map depicting the average, scaled expression of aberrant goblet markers across distinct groups of goblet cells. **E**. Ridge plot showing the distribution of the aberrant goblet cell signature per cell type and colored by the healthy (grey) and ulcerative colitis (red) data sets. **F**. Box plots showing the percentage of MUC2/MUC5AC double - and MUC2/MUC5AC/MKI67 triple positive cells per patient in the data sets from Smillie et al.^3^. P values denote the result of the Student’s t-test. **G**-**I**. Multiplex immunofluorescence images of a colon sample from an ulcerative colitis patient. **G**. Inset displays MUC5AC (yellow) positive ribbon. **H**. Inset displays MUC5AC (yellow) / MUC2 (purple) double-positive cells. **I**. Inset display examples of a 1) MUC5AC (yellow) / KI67 (magenta) positive cell and 2) MUC5AC positive cell with a signet ring-like morphology. UC: ulcerative colitis.

Next, we inferred the activity of key signaling pathways using decoupleR, an open-source package designed to infer biological activities from omics data by estimating enrichment scores for molecular processes^24^. This analysis revealed upregulation of the EGFR and MAPK signaling pathways in aberrant goblet cells, followed by WNT, PI3K, and ER, suggestive of reacquired proliferative capacity in UC (Figure 2C).

We subsequently asked whether aberrant goblet cells express markers unique to their cellular state. To this end, we performed differential expression analysis by comparing aberrant goblet cells with all other epithelial cells. This led to the identification of 153 genes upregulated in aberrant goblet cells. Using a clustering approach based on the average expression per cell type, we further refined a group of 10 genes uniquely upregulated in the aberrant goblet cell state: *MUC5AC, KLF2, KDELR3, ITLN1, HERPUD1, AGR2, REG4, PLXNA3, PHLDA2, OCIAD2* (‘aberrant goblet signature’; Suppl. Figure 3B). This gene signature was upregulated in UC compared to controls and, in particular, in the aberrant goblet cells identified by RNA velocity analysis (Figure 2D). Among the epithelial cells, the aberrant goblet signature mapped, as expected, to the goblet cell lineage, showing upregulation in UC in both mature and immature goblet cells as well as in secretory transit-amplifying (TAs) cells (Figure 2E). To identify cells with the highest activation of the aberrant goblet cell state, we established a threshold (aberrant goblet signature score > 1) and observed that, while rare in the data set from normal samples (0.01% of epithelium), they were represented in the UC data set (4.1% of epithelium) (Figure 2E).

To visualize the aberrant goblet cells *in situ*, we selected *MUC5AC* as a specific marker (Suppl. Figure 3C), along with *MUC2* as a control for the normal goblet cell lineage, and *KI67* as a marker for proliferative cells. In the scRNAseq data, *MUC2*^+^/*MUC5AC*^+^ double-positive cells were rarely observed in normal samples (0.01 ± 0.03% of epithelium). Instead, their abundance increased in UC colonic tissues (0.6 ± 1.06% of epithelium). The same was true for *MUC2*^+^/*MUC5AC*^+^/*MKI67*^+^ triple positive cells (<0.001% in normal vs. 0.17±0.34% in UC) (Figure 2F).

While absent in normal colon, IHC analysis revealed a strong and specific MUC5AC staining pattern in the colon tissues from UC patients in cells with a heterogeneous morphology ranging from signet ring-like cells to apparently normal goblet cells (Suppl. Figure 3D). Moreover, in specific patients, entire ribbons of MUCAC5^+^ cells were also observed (Figure 2G). Multiplex immunofluorescence (IF) analysis of MUC5AC and MUC2, and of MUC5AC and Ki67, confirmed their co-expression in the same cells, indicative of their goblet and proliferative characteristics (Figure 2H-I, Suppl. Figure 3E, Suppl. Video 1). Of note, a subset of MUC5AC^+^ cells identified in UC colonic tissues was earmarked by a distinctive signet-ring-like morphology (Figure 2H), similar to what was previously observed in signet ring cell carcinoma (SRCC)^25^.

In summary, these results indicate that goblet cells respond to inflammation by acquiring aberrant/immature gene profiles with proliferative capacity.

### Aberrant goblet cells as precursors of serrated colon tumorigenesis

To assess the potential relevance of the inflammation-driven reprogramming of goblet cells in colon tumorigenesis, we employed the newly developed aberrant goblet cell signature to mine the single-cell atlas from the Chen et al. study^4^ encompassing 122,811 cells from 121 pre-malignant polypoid lesions. First, to evaluate inter-polyp transcriptional heterogeneity, we performed UMAP dimension reduction after pseudo-bulk conversion of the epithelial cells in each polyp to project differences between serrated and conventional adenomas (Figure 3A). The aberrant goblet cell signature was strongly enriched in serrated lesions (Figure 3B). We next evaluated published expression signatures to capture the distinct types of RSC and CBC cells^15^. These signatures showed opposite associations with serrated lesions and conventional adenomas corresponding to enrichment of RSCs and CBCs, respectively (Figure 3C, D). Last, a recently developed human intestinal fetal signature^26^ differentially expressed in first-trimester progenitors compared to second-trimester colonocytes, was found to be upregulated in polyps relative to normal tissue, regardless of histological subtype (Suppl. Figure 4A).

**Figure 3.**
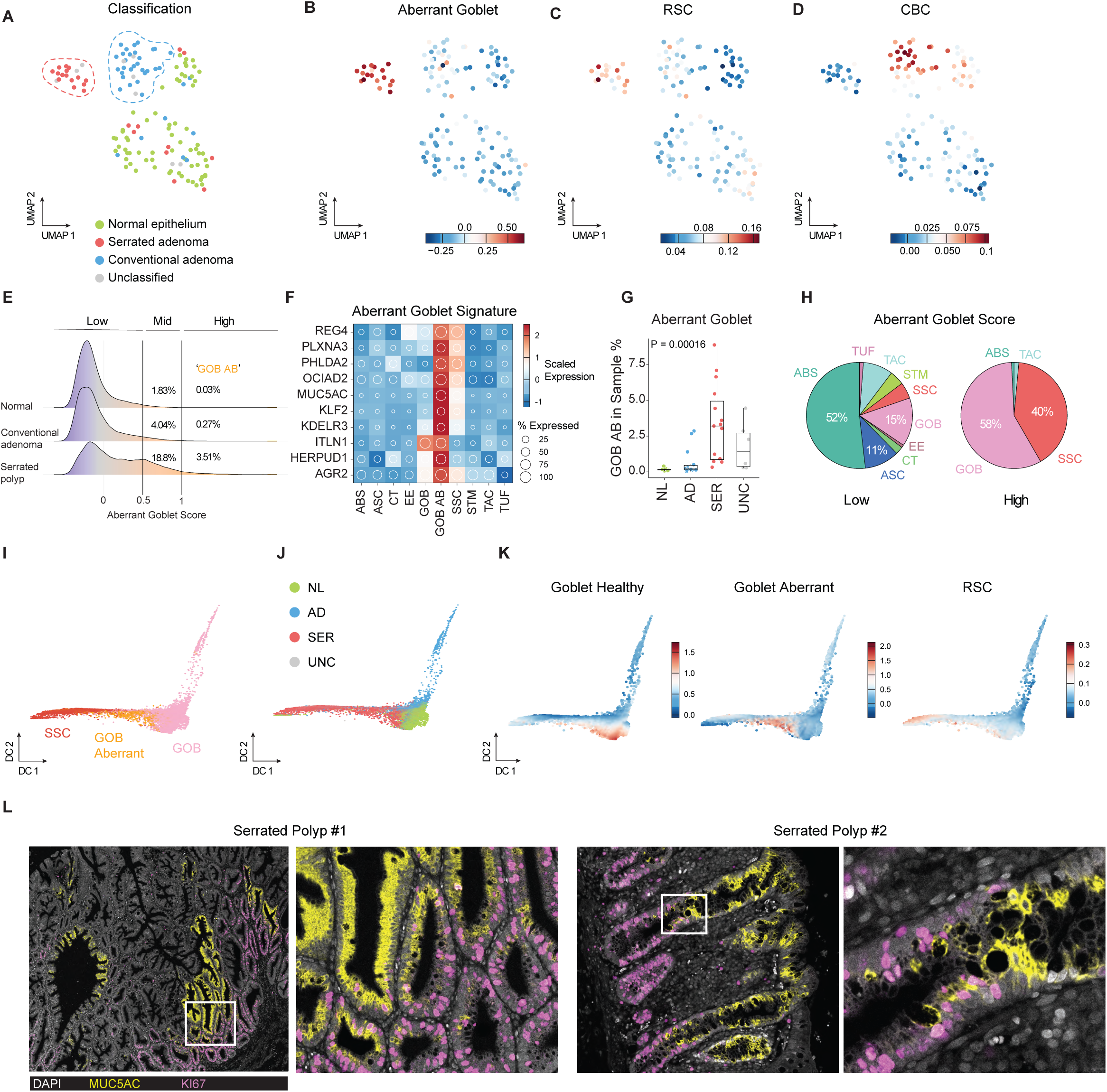
Aberrant goblet cells as precursor of serrated specific cancer cells. **A.** UMAP embedding showing the pseudo-bulk data of Chen et al.^4^. Dots represent individual polyps and are colored according to their pathological annotation. **B-D**. Feature plots showing the module scores for the (**B**) Aberrant goblet, (**C**) RSC, and (**D**) CBC signatures. **E**. Ridge plot displaying the distribution of the aberrant goblet cell signature across Chen et al.^4^ data set. Cells with a module score above 1 were classified as high/aberrant goblet. **F**. Heat map showing the average scaled expression of the aberrant goblet cell signature across the distinct cell types in Chen et al.^4^ with the addition of the aberrant goblet cell population (GOB AB). **G**. Box plot showing the percentage of aberrant goblet cells per polyp, stratified according to the pathological annotation. **H.** Pie chart showing the contribution of different cell types for cells with low (left) and high (right) association to the aberrant goblet signature. **I**. Diffusion component (DC) embedding of goblet (GOB, pink) and serrated specific cells (SSC, red). Aberrant goblet cells were colored in orange. **J**. DC embedding colored by polyp type. **K**. Feature plots of the DC embedding showing module scores for healthy, aberrant goblet cells, and RSC. **L**. Immunofluorescence imaging of serrated lesions. Serrated polyps were stained with antibodies against MUC5AC (yellow) and MKI67 (magenta). TUF: tuft cells, TAC: transit amplifying cells, STM: stem cells, SSC: serrated specific cell, GOB: goblet cells, EE: Enteroendocrine cells, CT: crypt top colonocyte (Best4^+^ cells), ASC: adenoma specific cells, ABS: absorptive cells. RSC: revival stem cell, CBC: crypt-base columnar cell.

To investigate in more detail the distribution of the aberrant goblet cell signature at the single cell level, we employed the same threshold as in our previous analysis of UC samples (aberrant goblet signature score > 1), and showed that only a rare (0.03%) cell subpopulation markedly expressed the aberrant goblet signature in the normal data set (Figure 3E). Compared to the healthy samples, their numbers increased in premalignant lesions (1.5%), and especially in serrated polyps (3.51%) when compared to conventional adenomas (0.27%).

As shown in Figure 3F, heatmap analysis confirmed that all 10 genes were expressed in the aberrant goblet cells indicating that the same gene expression module, identified in the UC context, earmarks a specific subpopulation in the polyp data set specifically enriched in serrated lesions (3.5% of cells) compared to conventional adenomas (0.27% of cells)(Figure 3G).

Further inspection of the cell annotations from Chen et al., matching the aberrant goblet cells indicated that the vast majority came from goblet (GOB: 58%) and serrated cells (SSCs: 40%)(Figure 3H). It should be pointed out that these annotations encompass cells from different sample types (i.e. normal and different types of polyps), and therefore represent a mixed population of healthy and premalignant cells. Given the large contribution of both GOB and SSC to the pool of aberrant goblet cells (98%), we addressed their relevance for colon tumorigenesis by investigating their potential reprogramming trajectory. To this aim, we utilized diffusion map embedding to separate cells in a low-dimensional space based on their probability to transit between states^27^ (see Methods)(Figure 3I-J). The diffusion embedding revealed that normal (green) goblet cells branched into either cells from serrated lesions (red) or conventional adenomas (blue)(Figure 3J). Interestingly, aberrant goblet cells (orange) mapped in between normal goblet (GOB) and serrated adenoma cells (SSCs), suggesting that these cells represent an intermediate state in the GOB to SSC transition. Notably, this reprogramming trajectory was exclusively observed in serrated lesions, accompanied by the activation of the revival stem cell (RSC) module (Figure 3K). In contrast, the branch connecting goblet cells to conventional adenomas displayed a closer resemblance to CBCs, rather than RSCs (Suppl. Figure 4B). Trajectory inference further indicated that goblet cells can reprogram into SSCs by activating an aberrant goblet cell state during the goblet-to-RSC transition (Suppl. Figure 4C). The aberrant goblet cell signature peaked at the GOB>SSC transition point, but returned to baseline levels once the full RSC state was established, suggesting a stepwise reprogramming process in serrated lesions (Suppl. Figure 4D).

IF analysis of sporadic serrated polyps confirmed the presence of extended ribbons of MUC5AC^+^ cells adjacent to fields of KI67^+^ cells with few double-positive cells located in the transition zone (Figure 3L, Suppl. Figure 4E).

In summary, these findings indicate that aberrant goblet cells are present in serrated lesions but not in conventional adenomas, and are likely to contribute to the onset of serrated colon tumorigenesis by transitioning into serrated-specific cells and activating the revival stem cell module.

### Tracing aberrant goblet cells in BRAF mutant colorectal cancer

In view of the high frequency of oncogenic *BRAF* mutations in serrated lesions^4,28^, we investigated whether *BRAF*-mutant CRCs encompass aberrant goblet cells. Initially, we assessed both the aberrant goblet cell and the RSC/CBC signatures in a panel of CRC cell lines^29,30^, ranking them based on the average expression of the aberrant goblet cell signature (Suppl. Figure 5A). Most of the cell lines earmarked by a high aberrant goblet score revealed increased expression of the RSC module, although this trend was not uniform across all lines (Suppl. Figure 5B). We selected the HT29 and LS174T cell lines for further *in vitro* validation based on their relatively increased expression of the aberrant goblet signature, and Caco-2 as a negative control with a low aberrant goblet score and RSC. Of note, both HT29 and LS174T are *BRAF*-mutant. Fluorescence-activated cell sorting (FACS) analysis of the selected cell lines revealed a subpopulation of MUC5AC^+^ cells in both LS174T and HT29, which was otherwise absent in Caco-2 (Suppl. Figure 5C). HT29 immunofluorescence analysis confirmed the presence of a cycling (KI67^+^) subpopulation of MUC5AC^+^ cells (Suppl. Figure 5D). Subcutaneous injections of HT29 and LS174T into NSG mice revealed a notable fraction of MUC5AC^+^ cells with patchy staining patterns (Suppl. Figure 5E). These experiments suggest that remnants of aberrant goblet cells remain detectable in *BRAF*-mutant CRCs.

Finally, we took advantage of whole-genome sequencing data from a cohort of n=165 sporadic colorectal cancers^5^ and computed cell-of-origin predictions based on the association between mutational spectra and epigenetic profiles of distinct cell lineages, as previously shown^2,31,32^ (Suppl. Figure 6). Stem cells were identified as the most dominant cell-of-origin among sporadic MSS cases, in confirmation of our previous observations^2^. Interestingly, the vast majority of MSI-H colon cancers were predicted to originate from a differentiated colonic lineages, in confirmation of a previous report with a distinct computational approach^33^, and in particular from goblet cells and enterocytes (Figure 4A). Compared to those predicted to originate from a stem cell, colon cancers arising from differentiated cells displayed distinct histological features, with a higher fraction of mucinous adenocarcinoma (stem: 5/58 [8.6%], differentiated: 19/64 [29.7%]). At the transcriptional level, lesions with a predicted stem origin showed increased association with the CBC signature, whereas tumors predicted to originate from goblet cells revealed increased expression of the RSC and aberrant goblet signature (Figure 4B). Subsequent evaluation of mutations in a panel of CRC driver genes indicated that tumors with a predicted differentiated cell-of-origin had a higher prevalence of mutations in *BRAF* (stem: 4/55 [7.3%]; differentiated: 25/61 [41.0%]) and *RNF43* (stem: 0/55 [0%]; differentiated: 24/61 [39.3%]), and lower prevalence of *APC* (stem: 47/55 [85.5%]; differentiated: 37/61 [60.7%]) and *TP53* mutations (stem: 29/55 [52.7%]; differentiated: 20/61 [32.8%]), when compared to those predicted to originate from stem cells (Figure 4C). Thus, the selection of key colon cancer driver genes, may at least partly depend on the tumor cell-of-origin.

**Figure 4.**
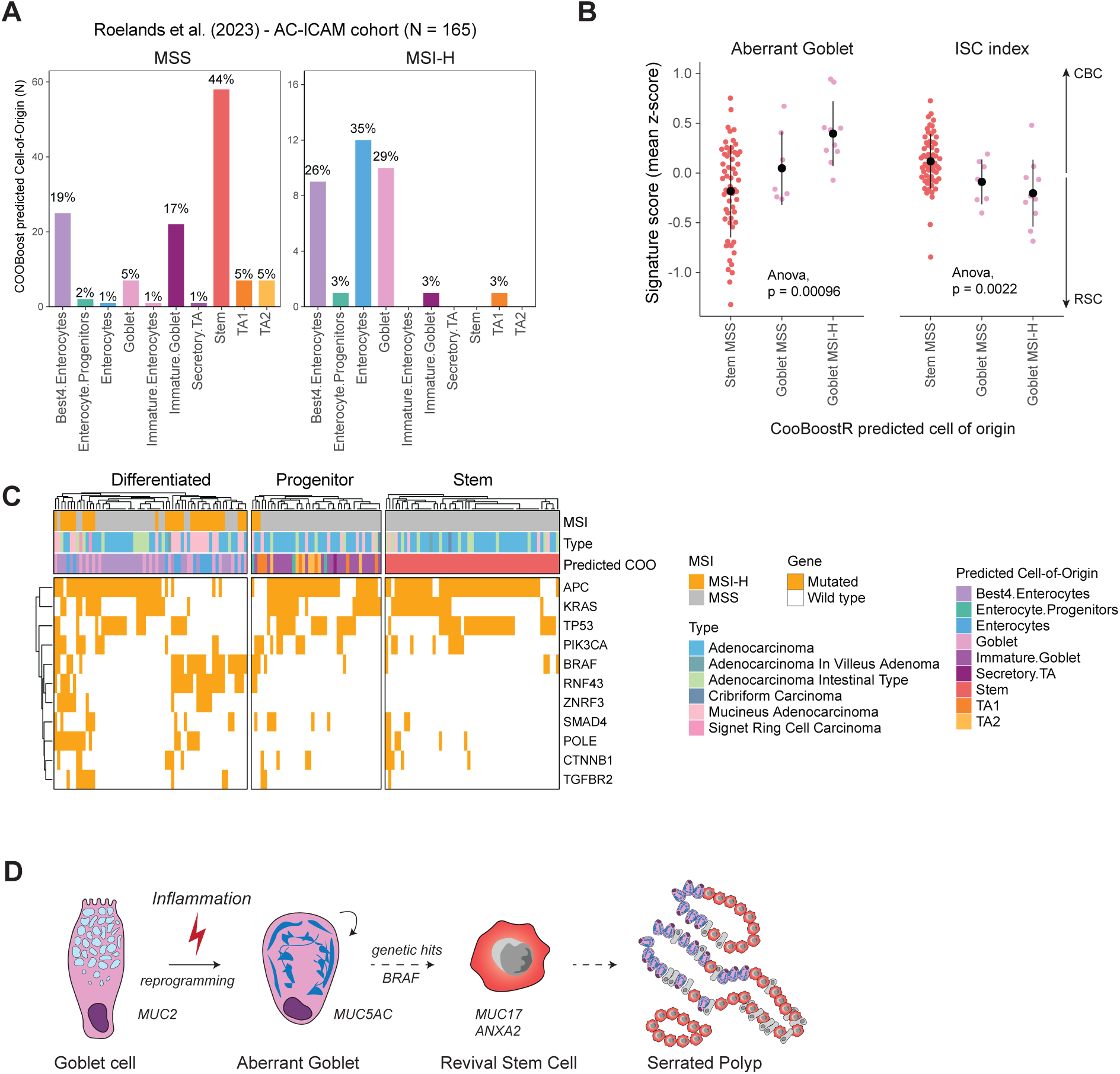
Traces of aberrant goblet cells in BRAF mutant colon cancer. **A.** Barplot showing distribution of tumors from the AC-ICAM cohort stratified by MSS/MSI-H and the predicted cell-of-origin using CooBoostR. **B.** Violin stratified by case with a predicted Stem or Goblet cell-of-origin. Plots showing the distribution of revival stem cell (RSC), the crypt-base columnar cell (CBC), and Aberrant Goblet cell signatures. **C**. Heatmap showing driver gene mutation status (white: wildtype, orange: mutated) across colon cancers stratified by their predicted cell-of-origin. **D**. Schematic illustration of proposed model. Goblet cells reprogram in inflammatory context to MUC5AC aberrant goblet cells. Upon genetic hits such as oncogenic BRAF mutation, these cells may progress to revival stem cells (RSCs) in serrated lesions.

These observations, together with the role of aberrant goblet cells as precursors of serrated (and often *BRAF*-mutant) adenomas, support a scenario where the high-grade and chronic inflammation features of ulcerative colitis results in the reprogramming of goblet cells leading to the onset of premalignant lesions with distinct serrated histology and somatic (e.g. in *BRAF* and *RNF3*) mutations, eventually progressing to RSC-enriched aggressive adenocarcinomas with poor prognosis and overall survival (Figure 4D)^15^.

## Discussion

Inflammation is nowadays regarded as the single phenomenon that most contributes to the medical burden of industrialized societies, being associated with almost every major non-communicable disease^34,35^. Nonetheless, the molecular and cellular mechanisms underlying inflammation-driven disease initiation, rather than progression, are still poorly understood. From this perspective, patients suffering from inflammatory bowel diseases, such as Crohn’s disease (CD) and ulcerative colitis (UC), characterized by a chronic and relapsing immune-mediated inflammation of the digestive tract, represent unique models to study the short- and long-term consequences on tissue homeostasis and colon cancer onset and progression.

In this study, we characterized the impact of chronic inflammation of the colon on the epigenetic reprogramming of the crypt of Lieberkühn, by taking advantage of extensive and readily available single-cell RNAseq data from UC patients, and of a novel computational RNA velocity-based approach. Our findings reveal that UC induces lineage-specific reprogramming of the colonic epithelium. We specifically focused on the goblet cell lineage in view of our previous observations, according to which these secretory cells represent the most common non-stem origin of colon cancer, not only in the context of IBD but also among sporadic microsatellite-stable^2^ and, as shown here, in MSI-H cases.

We identified an aberrant goblet cell gene signature that is selectively activated in colonic biopsies from ulcerative colitis patients. Expression of the *MUC5AC* gene encoding for a gastric mucin, earmarked the aberrant goblet cell signature. Of note, *MUC5AC* is not only frequently detected in serrated polyps^4^ but also in UC-associated colon carcinomas^36^. In mouse models of dextran sulfate sodium (DSS)-induced colitis, expression of *Muc5ac* was shown to protect the colonic barrier^37^. More recently, *MUC5AC* expression was identified as a specific marker of the inflammatory epithelial cells (INFLAREs) located at the crypt top (surface foveolar cells) of metaplastic glands in Crohn’s disease, thought to result from stem cell differentiation in the context of intestinal injury and to underlie pyloric metaplasia in patients with Crohn’s disease^38^.

Our analyses indicate that aberrant goblet cells are likely to directly arise from differentiated goblet cells. However, we cannot exclude that a fraction of aberrant goblet cells can emerge from aberrant differentiation of intestinal stem/progenitor cells or through an RSC intermediate state, as also observed in mouse models^2^. Hence, the pool of *MUC5AC-*positive cells may represent a complex mixture of cells derived from distinct lineages.

While some of the genes activated in aberrant goblet cells overlap with previously reported fetal-like state^15^, we did not observe reversion to RSCs in colitis. However, a subset of tumor cells from pre-malignant serrated adenomas is earmarked by the RSC signature, likely linked to MAPK activation triggered by *BRAF* oncogenic activation^39^. Trajectory analysis indicated that aberrant goblet cells represent an intermediate state in the goblet-to-RSC conversion. Hence, we propose that inflammation triggers reprogramming of goblet cells toward aberrant goblet cells, which, in turn, may be primed for RSC-conversion upon acquisition of genetic hits. Complete RSC conversion likely requires additional reprogramming attained by oncogenic hits or by tissue injury, as shown by γ-irradiation in mice^14^.

Of note, we observed a subset of MUC5AC^+^ cells with a distinctive signet-ring-like morphology in both colonic tissues from UC patients and in sporadic serrated adenomas. Given their prominent presence in signet ring cell carcinoma (SRCC), an aggressive subtype of CRC^25^, it remains to be determined whether their presence in pre-malignant epithelial tissues serves as a marker of a differentiated cell-of-origin and of poor prognosis.

The striking predominance of sporadic MSI-H colon cancers predicted to originate from a committed cell lineage (>90%, mainly enterocytes and goblet cells), sharply contrasts the situation among MSS cases (44% predicted to originate from stem cells). Whereas metaflammation due to Western-style dietary habits^40^ represents a likely cause for the non-stem origin in ∼40% of the sporadic MSS-CRCs, the cause of inflammation among MSI-H cases might be different. Of note, the preferential inactivation of the *TGFBR2* gene (encompassing an A10 repeat in its coding sequence) in the context of MSI-H (both in the normal colon^41^ and in CRC^42^) and its role in inflammation^43,44^, might indicate that distinct inflammatory insults result in the epigenetic reprogramming of distinct lineages.

### Limitations of the study

While RNA velocity represents a powerful framework to study cellular plasticity, it is a modeling approach with some limitations that cannot be readily applied to any data set without careful evaluation of the results^16^. Given the sparsity of single-cell expression data and the inherent heterogeneity across bio-specimens, the method necessitates a balance between minimizing noise and technical effects while retaining the biological signal. Additionally, the modeling results depend on dimensionality reduction and data integration, which can influence the outcome^45^. For datasets with heterogeneous samples and multiple cell lineages, it is essential to carefully inspect gene phase portraits, as the default gene selection in our models included a large portion of noisy genes. To address this, we conducted thorough evaluations of model results on the healthy dataset before any comparisons with the ulcerative colitis data.

Collectively, our study contributes to our understanding of inflammation-driven cellular reprogramming and its relevance for tumor onset. Of note, in view of the chronic “metaflammation” caused by Western-style dietary habits^40,46^, i.e. the main etiologic colon cancer risk factor, and the striking frequency at which sporadic, non-IBD, MSS and MSI-H colon cancers are predicted to originate from differentiated lineages^2^, it is plausible to think that inflammation, driven either by unhealthy dietary habits or by somatic inactivation of target genes (e.g. *TGFBR2*), underlies lineage-specific epigenetic reprogramming and tumor onset in a large proportion of colon cancers in human. Future studies leveraging genetically engineered mouse models and advanced organoid systems will likely help unravel the complex cellular states induced by etiological factors such as diet and inflammation, which underlie the onset of (serrated) adenomas and their progression towards malignancy.

## Methods

### Preprocessing of ulcerative colitis single cell RNA sequencing data

Raw data from Smillie et al.^3^ was retrieved from TerraBio. Fastq files were aligned to the human reference genome GRCh38-3.0.0 with Cellranger. Loom files, containing raw spliced and un-spliced counts, were produced by *velocyto run10x*. Original annotations and metadata from the original paper^3^ were added post-pre-processing. The full dataset consists of 17 ulcerative colitis (UC) patients and 12 healthy individuals, partitioned into three major cell types: immune, stromal, and epithelial. We restricted our analysis to the epithelial cells.

### Batch correction and dimension reduction

Batch effects were corrected using the Seurat^47^ integration method with the functions *FindIntegrationAnchors* and *IntegrateData* based on default parameters. These functions were implemented using canonical correlation analysis (CCA) to identify matching cell pairs across datasets called anchors. Uniform Manifold Approximation and Projection (UMAP) was employed using the top 50 principal components with the Seurat function *RunUMAP* with 200 neighbours and a minimum distance of 0.3.

### Trajectory inference

Trajectory inference was performed with Monocle3^48^. The integrated Seurat object was converted to Monocle object with SeuratWrappers::as.cell_data_set. Subsequently, cluster cells and learn graph were computed with default parameters. The order cells function was utilized with the stem cells as appointed root neighbourhood. Subsequent trajectories were visualized with the plot cells function, where cells were coloured by the pseudotime value.

### Differential abundance

Abundance analysis was performed with MiloR^19^. The kNN-graph was constructed using the integrated_nn graph from Seurat (k = 20 nearest neighbours). Next, the *makeNhoods* function was utilized with prop = 0.05 and d = 30 dimensions based on reduced dims = “PCA”. The number of cells in each neighbourhood was quantified per patient using countCells function. Healthy vs. UC comparison was done with *testNhoods* (fdr.weighting = “neighbour-distance”). Neighbourhoods with less than 70% overlap to one annotation were annotated as mixed and excluded from the analysis.

### Single cell Entropy

Entropy values were computed using SCENT^23^. The net13Jun12 data was used as protein-protein interaction network. Entropy values were calculated on all epithelial cells with the *CompSRana* function.

### RNA velocity analysis

RNA velocity analysis was performed with scVelo 2.4^18^. Cells with less than 20 shared spliced and unspliced counts were excluded. Next, data were normalized and log-transformed following the default parameters of the scVelo implementation. We calculated the first and second-order moments with the sklearn method on neighbourhoods of 200 cells using the dynamical modelling estimation to compute velocity vectors. To optimize the input velocity genes, we visually inspected the velocity genes in the spliced vs. unspliced scatter plots, which revealed a considerable fraction of genes with noisy phase portraits. To identify the noisy genes, we clustered the transposed count matrix based on UMAP embedding with k-means (k = 3). Subsequently, after evaluation of the log2(counts) and manual inspection of phase-portraits, one cluster was removed from the input velocity genes. The remaining clusters encompassed N = 834 genes, and were used for the subsequent analysis. Both the healthy and ulcerative colitis data set were processed independently using the same settings and velocity genes as input.

### Definition of aberrant cells

Single cell velocity vectors were projected on the UMAP embedding. For each annotation, the average direction was computed in the healthy data set based on the angle (theta). Cells with an angle two standard deviations away from the average were annotated as aberrant cells.

### Gene signatures and pathway analysis

The signatures of healthy cell lineages (e.g. Healthy Goblet, Healthy Immature Goblet) were obtained from Smillie et al. ^3^ supplementary information. The signatures for RSC and CBC stem profiles were retrieved from Gil Vazquez et al.^15^ Signatures were evaluated with the *AddModuleScore* function in Seurat. Pathway activity scores were computed with decoupleR based on the PROGENy model gene weights (top = 500). The *run_ulm* function was used to compute pathway activity scores. After scaling with *ScaleData* function in Seurat, activity was visualized as mean per cell group using the ComplexHeatmap package.

### Aberrant goblet cell signature

To identify markers for the aberrant goblet cells, a differential expression analysis was performed by comparing the aberrant goblet cells against all other epithelial cells using the *FindMarkers* function (logfc.threshold = 0.25, only.pos = T, test.use = “wilcox”). Subsequently, the gene list was filtered using a k-means clustering analysis (k = 4) based on the average scaled expression of healthy, UC and UC aberrant cell groups. The cluster specifically expressed in the aberrant goblet cells was selected. Only genes with p_adj < 0.01 and pct.1 > 0.1 were selected as aberrant goblet cell signature. The resulting gene list consisted of 10 genes: *MUC5AC, KLF2, PLXNA3, REG4, KDELR3, ITLN1, PHLDA2, AGR2, HERPUD1, OCIAD2*.

### Processing of polyp single cell RNA sequencing data

The epithelial DIS and VAL data sets from Chen et al. ^4^ were retrieved from the cell-x-gene database. The h5ad files were converted to h5seurat files and merged as one Seurat object. The signatures as described above were calculated in the data set using the *AddModuleScore* function from Seurat. Pseudobulk analysis was performed with the *AggregateExpression* function based on HTAN.Parent.Data.File.ID and Sample Classification columns. Subsequently, the pseudobulk data was processed with *NormalizeData()*, *FindVariableFeatures* (nfeatures = 3000), *ScaleData(), RunPCA()* and *RunUMAP* (dims = 1:20) functions. Signatures were visualized with FeaturePlots in the pseudobulk data set, and as Ridgeplots in the single cell data set.

### Pseudo-time analysis of goblet cells and serrated-specific cells

A subset was created containing goblet (GOB) and serrated-specific cells (SSC). To compute diffusion map of cells, the ‘Diffusion_Map’ function from the destiny package^27^ was employed with default parameters based on the n_pcs = 30 principal components. Genes with fewer than 100 counts were filtered from this analysis. To create pseudotime ordering of the cells, the Diffusion Pseudo Time ‘DPT’ function was employed. The cell with the highest value of the ‘Healthy Goblet’ signature was chosen as the root cell. The diffusion map was visualized with the SCpubr package. Cells from the serrated adenomas were further selected for visualization with a heat map. Cells were ordered according to their respective pseudotime values, and scaled expression was smoothened based on a rolling average (window = 300, by = 50) of the top 250 variable-expressed genes. The heat map was visualized with the Complex_Heatmap package.

### Evaluation of aberrant goblet cell signature in CRC cell lines

Data from Zhu et al.^30^ and Kinker et al.^29^ was processed by Seurat. Only colorectal cancer cell lines were selected, and the two studies were merged. Subsequently, signatures were evaluated with the Add_Module_Score function. Cell lines were ranked based on the expression of the Aberrant Goblet signature; signature distributions were shown with ridge plots.

### BRAF mutations and CRC with predicted goblet-origin

Whole-genome sequencing data from Roelands et al.^5^ were processed by counting the number of mutations across 1 Mb genomic bins, similar to our previous study^2^. Data from Becker et al.^49^ was aggregated using the same genomic bins, and counts were averaged by cell type using the Aggregate_Expression function (slot = ‘counts’). Cell-of-origin predictions were done using CooBoostR with parameters mEta = 0.3, mdepth = 6 ^32^. Samples were stratified according to their predicted cell-of-origin, and data from cBioPortal (id=coad_silu_2022) were crossed to evaluate the effect of cell-of-origin on histology and BRAF mutation.

### Immunohistochemistry analysis of MUC5AC

Chromogenic double labelling was performed on 4-µm thick whole slide sections from FFPE tissue blocks, on a validated and accredited automated slide stainer (Benchmark ULTRA System, VENTANA Medical Systems, Tucson, AZ, USA), according to the manufacturer’s instructions. Briefly, following deparaffinization and heat-induced antigen retrieval for 40 minutes at 97°C, the tissue samples were incubated with MUC5AC antibody (V2198 NSJ Bioreagents, 45M1, 1:700), followed by Ultraview detection (#760-500, Ventana). Counterstain was done by hematoxylin II for 12 min and a blue colouring reagent for 8 min. Each tissue slide contained a fragment of FFPE stomach as an on-slide positive control for MUC5AC.

### Cell culture

Human colon cancer cell lines were cultured in DMEM medium (#11965092, Thermo Fisher Scientific), supplemented with 10% heat-inactivated fetal calf serum, 1% penicillin/streptomycin (penicillin: 100 U/mL, streptomycin: 100 µg/mL; #15140122 Thermo Fisher Scientific) in humidified atmosphere at 37°C and 5% CO2. Cell lines were tested for *Mycoplasma pulmonis* and an additional panel of viruses by the RAP/MAP test (QM diagnostics). The results of these tests were negative.

### Fluorescence activated cell sorting

Fixed cells were permeabilized with 0.1% Triton for 20 min at room temperature. Following blocking with 5% FCS in HBSS, cells were stained with MUC5AC antibody (V2198 NSJ Bioreagents, 45M1, 1:500) or with mouse IgG (1:500) for 1 hour at 4°C. After two washes with 5% FCS in HBSS, cells were stained for 30 min on ice with Alexa anti-mouse 488 (A32723, 1:500). Cells were counterstained with DAPI to remove doublets, and FACS analysis was performed using FACSAria III (BD Biosciences). DAPI was detected using a 405 nm laser and 450/40 band-pass filter. Alexa-488 was detected using a 488 nm laser and 502 long-pass + 530/30 band-pass filters. Analysis of FACS data was performed with FACSDiva (v.8.0.1).

### HT29 IF analysis

HT29 cells were cultured on glass coverslips for 4 days and fixed with 4% PFA for 1 hour at room temperature. After 2 washes with PBS, cells were permeabilized with 0.1% Triton for 20 min at room temperature. Following 1h blocking with 5% milk in PBS, cells were stained with primary antibodies against MUC5AC (V2198 NSJ Bioreagents, 45M1, 1:700) and KI67 (MIB-1 antibody, 0.4 µg/mL, #790-4286; Ventana) overnight at 4°C. After two washes with PBS-Tween (PBS-T), samples were stained with secondary antibodies Alexa anti-mouse 488 (A32723, 1:500) and Alexa anti-rabbit 568 (A11011, 1:500). Cells were counterstained with DAPI (Sigma) and, where indicated, with F-actin (#SC001, Spirochrome) for 30 min. at room temperature and washed in PBS-T. Tissues were mounted in VECTASHIELD HardSet Antifade Mounting Medium (Vector Labs). Images were taken with the Stellaris 5 confocal microscope (Leica) and images were processed with ImageJ.

### Animal experiments

All animal procedures were approved by the Dutch Animal Experimental Committee and adhered to the Code of Practice for Animal Experiments in Cancer Research, as outlined by the Netherlands Inspectorate for Health Protections, Commodities and Veterinary Public health. Animals were bred and housed in the Erasmus MC animal facility (EDC) under specific pathogen-free (SPF) conditions. Subcutaneous injections were performed in 6- to 8-week-old male and female NSG (NOD.Cg-Prkdcscid Il2rgtm1Wjl/SzJ) mice. Briefly, 1×10^6^ cells were resuspended in 200 μl PBS and injected subcutaneously into the flanks of each mouse (4 injections per animal). Mice were euthanized once the tumor size reached the predetermined humane endpoint. Primary tumors were then excised, fixed in 4% paraformaldehyde (PFA), embedded in paraffin, and sectioned for subsequent immunohistochemistry analysis.

### Statistics and Reproducibility

Statistical tests were performed in R version 4.3.2. Statistical tests were performed with the ggpubr and rstatix packages. Representative micrographs come from at least N = 3 biologically independent samples, and display results that were observed consistently throughout a series of independent experiments.

## Data availability

All data relevant to this study are publicly available. Human scRNAseq data from Ulcerative Colitis patients (Smillie et al. ^3^) were downloaded from the Broad DUOS platform after a data transfer agreement. Single cell data from pre-malignant polyps have been reported in Chen et al.^4^; both discovery (DIS) and validation (VAL) epithelial data sets were obtained from the cell-x-gene platform (cellxgene.cziscience.com). Single cell profiling data of CRC cell lines were obtained from Zhu et al.^30^ using the China National GeneBank DataBase Sequence Archive (CNSA: identifier CNP0003658) and from Kinker et al.^29^ using the Gene Expression Omnibus (GEO: identifier GSE157220). Single-cell ATAC data from Becker et al.^49^ was obtained from GEO (GSE201349). Processed data from Roelands et al.^5^ was obtained from cBioPortal (id=coad_silu_2022).

## Code availability

The scripts that were used to run the analysis presented in this research are uploaded to GitHub (github.com/mpverhagen/AberrantGoblet).

## Patient-derived samples

Archival diagnostic colon specimens from serrated adenoma and IBD patients were used in accordance with the Dutch Code of Conduct for Health Research, issued by COREON, which provides guidance for the responsible use of human tissue and associated health data in medical research. Only residual material remaining after completion of routine diagnostic procedures was used. The use of tissue did not interfere with diagnostic assessment or future patient care.

## Acknowledgements

We thank all members of the Fodde laboratory for their input. We thank the PARTS research facility for help with tissue processing and immunohistochemistry experiments. This study was financially supported by the Dutch Cancer Society (KWF; project no. 11407).

## Author contributions

M.P.V., R.J., and A.S. designed and performed experiments. W.S. processed raw sequencing data sets. M.P.V. and E.M. analyzed sequencing data sets, and E.M. optimized and performed RNA velocity analysis. K.L., A.N. and T.P.P.v.d.B. performed immunohistochemistry experiments. D.S. and M.D. were responsible for pathological examination and IBD patient selection. H.S. Processed the AC-ICAM WGS data and generate the mutation count matrix. E.I.A. processed the AC-ICAM whole genome sequencing data and J.R. and W.H. contributed to interpretation of cell-of-origin predictions. M.P.V. and R.F. wrote the manuscript. R.F. supervised the study and was responsible for the concept and design of the study.

## Supplementary Figures

**Supplementary Figure 1.**
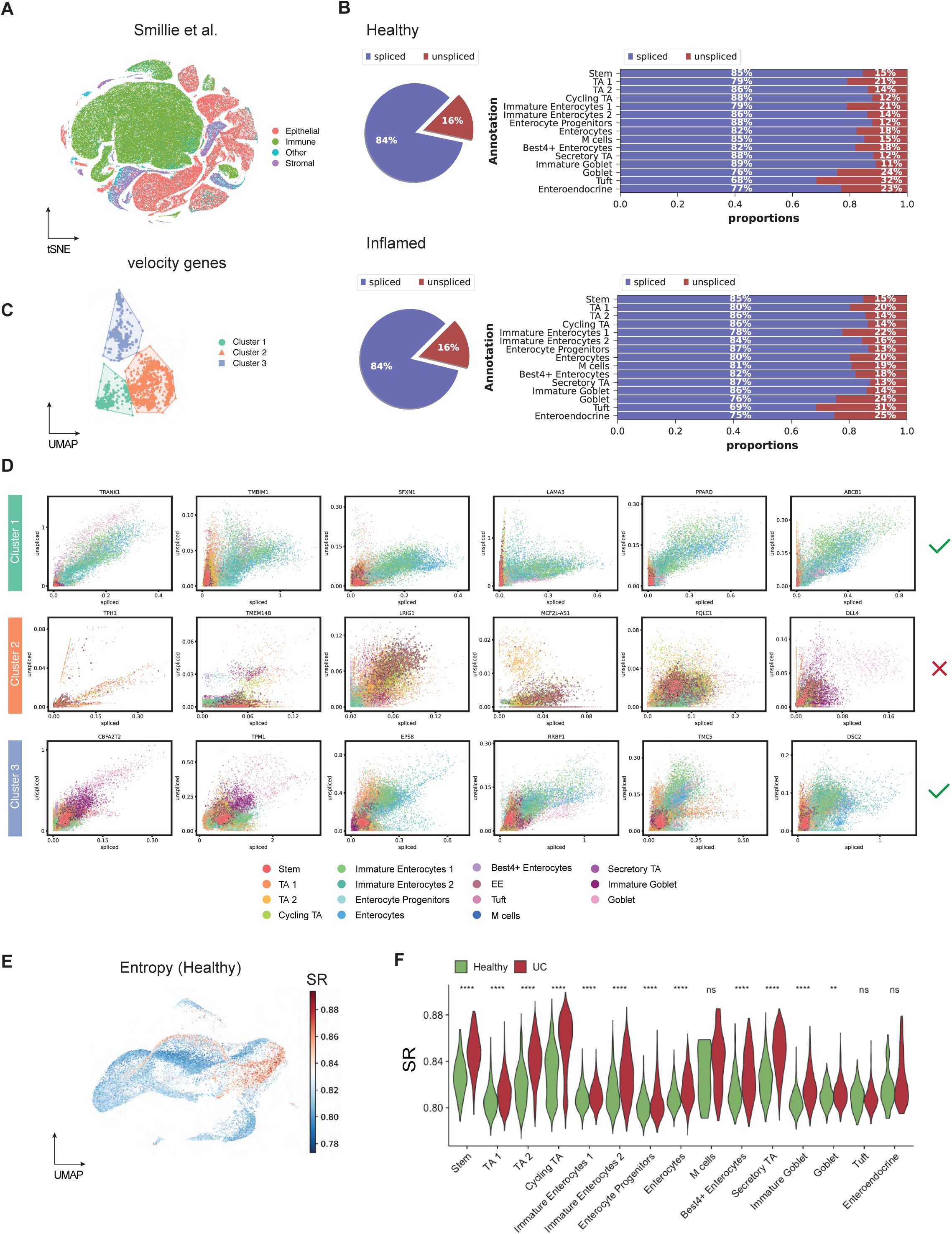
Optimization of the RNA velocity model. **A.** t-distributed stochastic neighbor embedding (tSNE) of the complete data set from Smillie et al. ^3^ **B**. Fraction of splice and unspliced reads per cell type and disease label. **C**. UMAP embedding of the RNA velocity genes based on the transposed count matrix. Cells were clustered by k-means (k = 3). **D**. Examples of phase portraits for the different k-means clusters. Cluster 2 was removed from the list of input genes for subsequent analyses **E**. Feature plot showing the signaling entropy [SR] values in the healthy data set. **F**. Violin plots comparing the SR values between healthy and UC data sets across the distinct cell types.

**Supplementary Figure 2.**
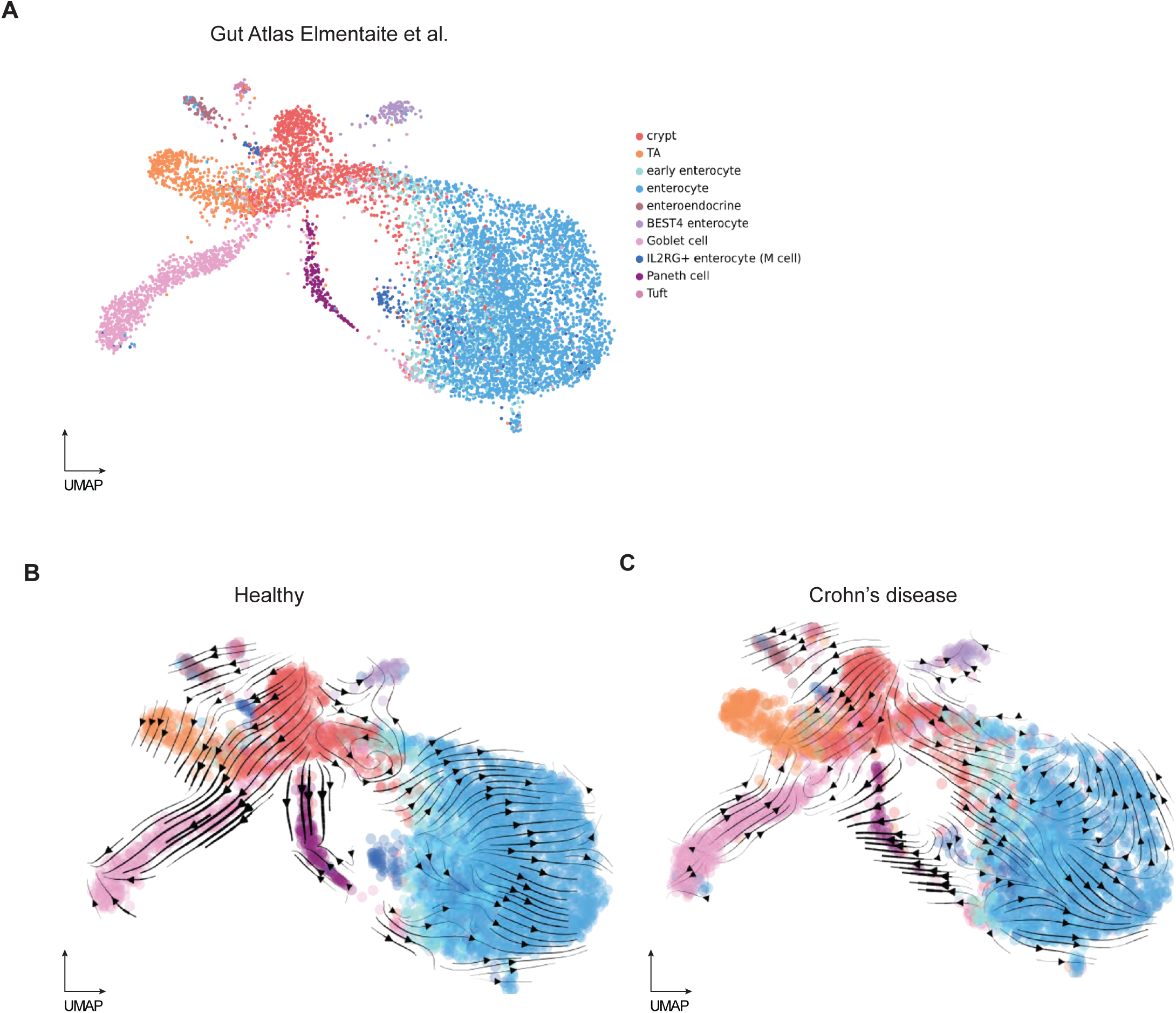
Application of velocity gene panel on an independent data set. **A.** Dimension reduction plot of the gut atlas described in Elmentaite et al.^22^**. B-C.** scVelo stream plots showing the predicted dynamics in the (**B**) healthy and (**C**) Crohn’s data set.

**Supplementary Figure 3.**
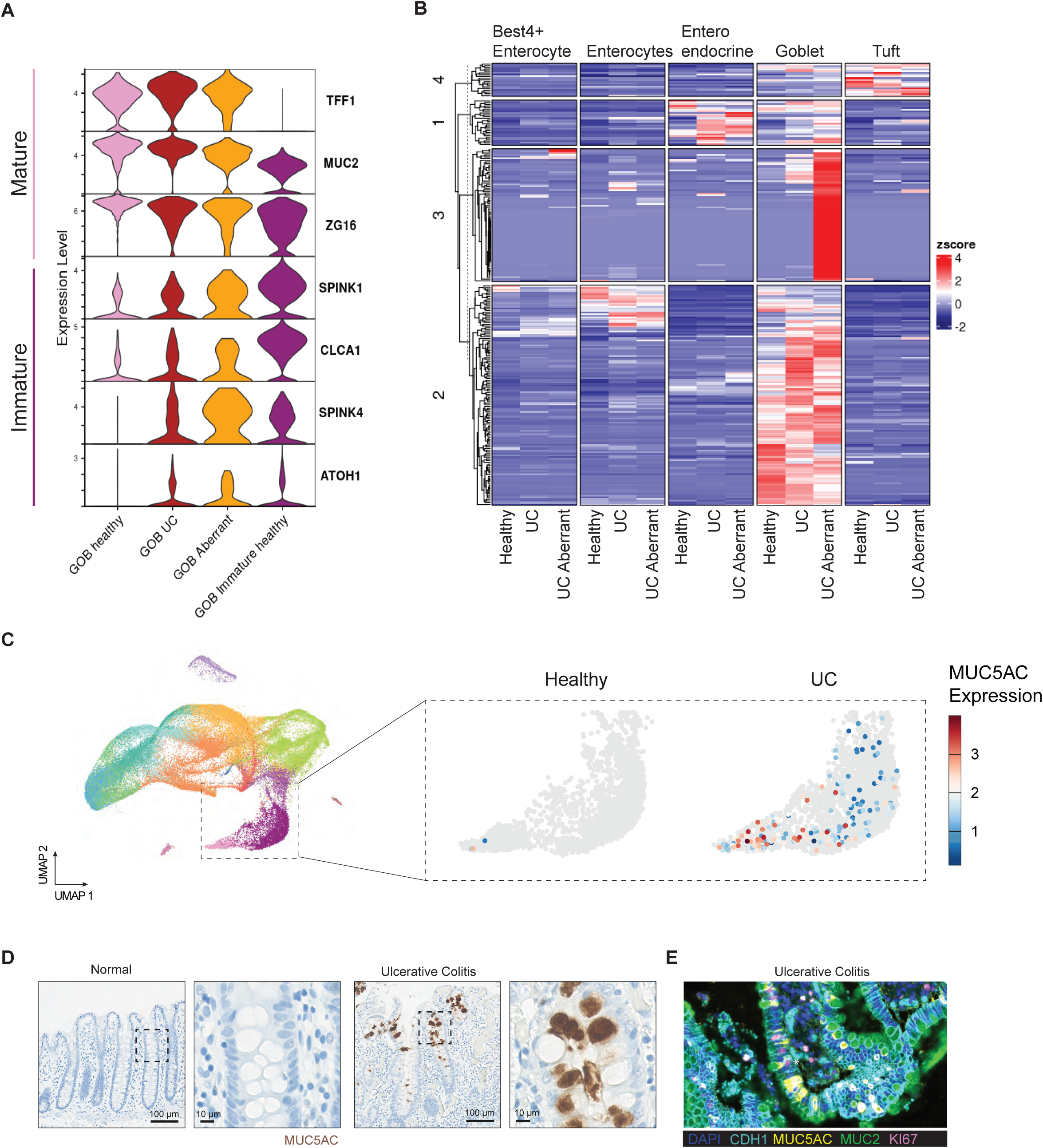
Identification of aberrant goblet cell markers. **A.** Stacked violin plot showing the expression of mature and immature goblet markers across the distinct groups of goblet cells. **B**. Heat map with k-means clustering showing the average expression of aberrant goblet cell markers throughout distinct cell groups. Cluster 3 depicts markers specific to the aberrant goblet cell group. **C**. Dimension reduction plot (left) and feature plot (right) showing the expression pattern of MUC5AC in goblet cells. **D**. Immunohistochemistry images based on MUC5AC staining in colon from normal and ulcerative colitis samples. **E**. Multiplex-immunofluorescence image of colon sample from an ulcerative colitis patient. MUC5AC/MUC2/KI67 triple-positive cells highlighted with a white asterisk and imaged with high magnification in Suppl. Video 1.

**Supplementary Figure 4.**
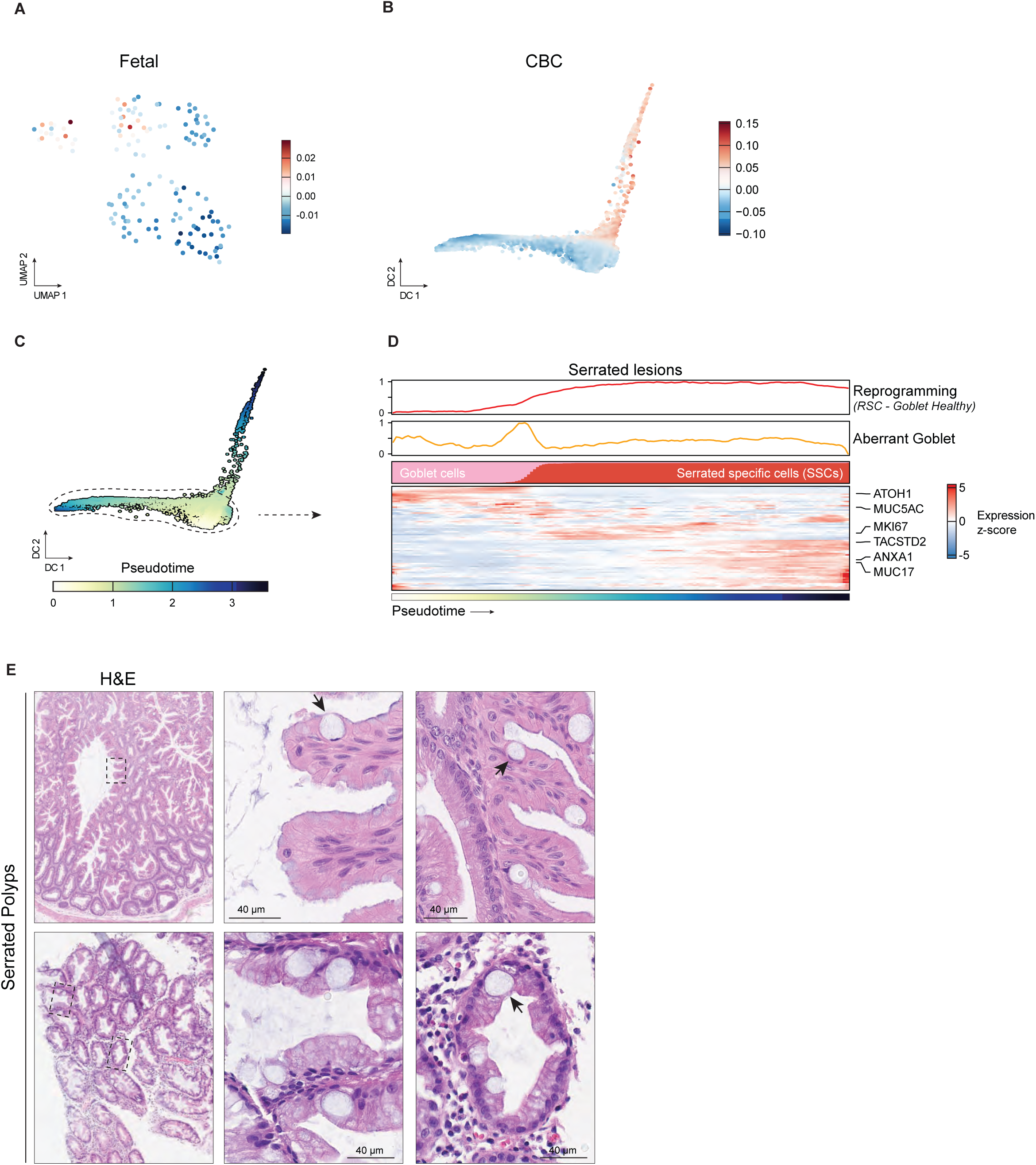
Aberrant goblet cells in serrated lesions. **A.** UMAP dimension reduction plot showing the fetal signature from Moorman et al.^26^ in the Chen et al. data set^4^. **B.** Diffusion component plot showing the crypt-base columnar (CBC) cell signature in the subset of Goblet (GOB) and Serrated-specific cells (SSCs) in Chen et al.^4^. **C**. Diffusion component embedding colored by pseudo-time ordering using destiny. **D**. Modeling the transition of goblet cells to serrated specific cells in serrated lesions. Heat map depicting pseudo-time ordered cells with smoothened expression and module scores. **E**. HE staining of consecutive slides. Examples of cells with signet-ring cell (SRC) morphology are highlighted with black arrows.

**Supplementary Figure 5.**
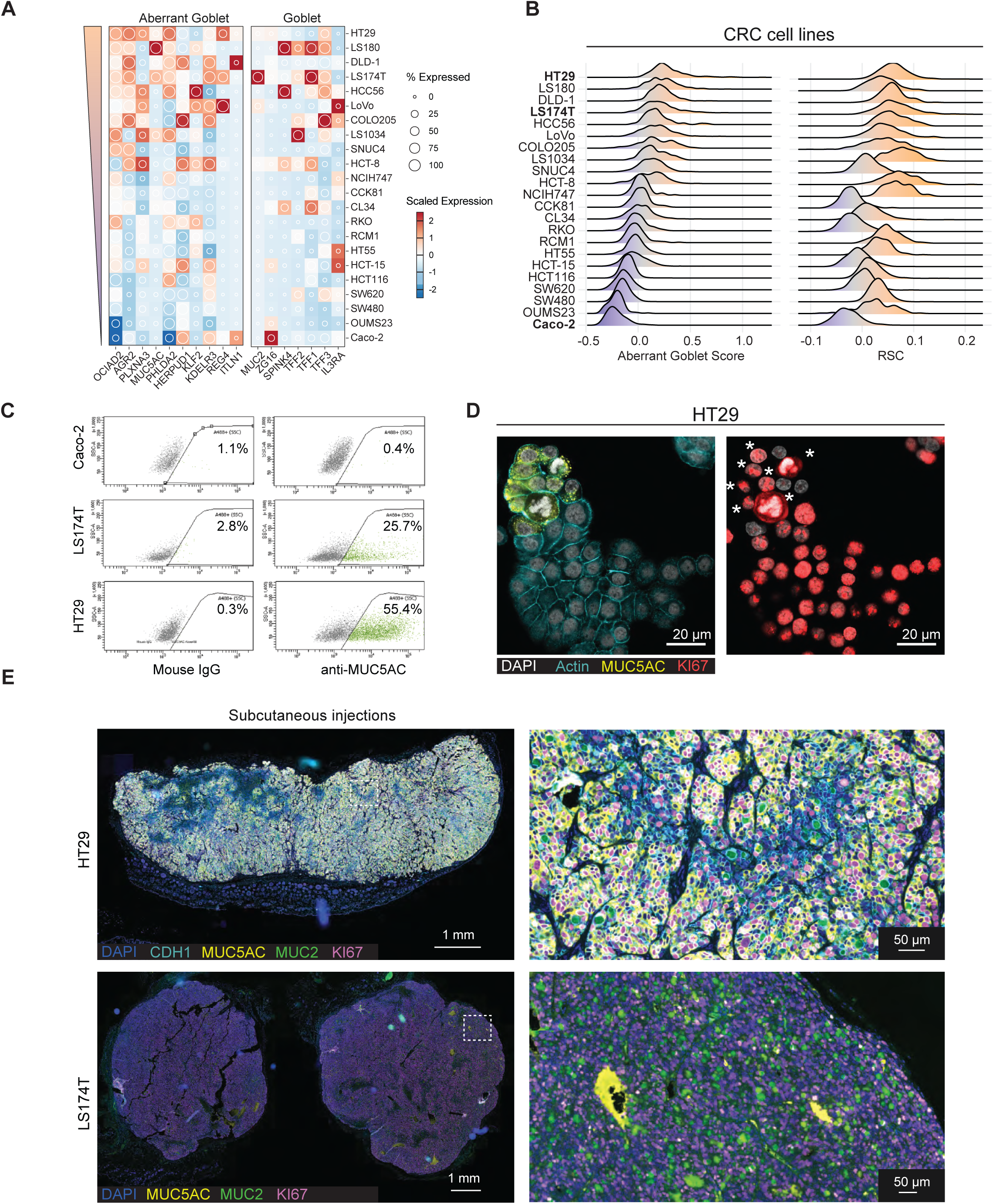
Traces of aberrant goblet cells in CRC cell lines. **A**. Heatmap showing average scaled expression of aberrant (left) and canonical (right) goblet markers across a panel of CRC lines. Data was aggregated from Kinker et al.^29^ and Zhu et al.^30^ and cell lines were ranked according to their association with the aberrant goblet cell signature. **B**. Ridge plot showing the distribution of the aberrant goblet signature (left) and the RSC signature (right) across the cohort of CRC cell lines. Cell lines were ordered based on the average of the aberrant goblet cell signature. HT29, LS174T and Caco2 (bold) were selected for follow-up experiments. **C**. Flow cytometry analysis of Caco-2, LS174T and HT29. Scatter plots highlight the percentage of MUC5AC-positive (green) cells. **D**. Multiplex immunofluorescence images of HT29 colon cancer cells. A subset of cells is positive for MUC5AC (yellow). MUC5AC/KI67-positive (red) double cells were marked with white asterisks. **E**. Multiplex immunofluorescence images of subcutaneous tumors from HT29 (top) and LS174T (bottom). Images represent four different tumors per cell line.

**Supplementary Figure 6.**
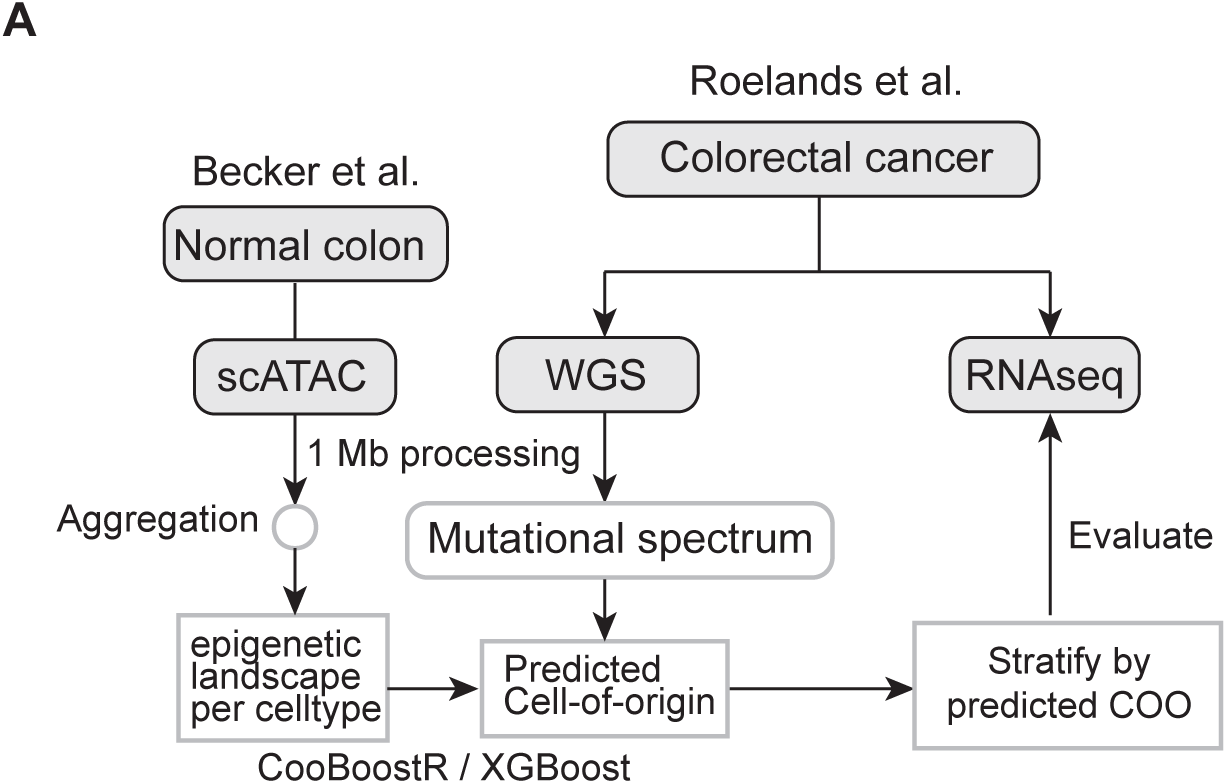
Schematic computational workflow for cell-of-origin predictions. Datasets from Becker et al.^49^ and Roelands et al.^5^ were processed using 1 Mb genomic bins and cell-of-origin predictions were performed with CooBoostR^32^.

